# Intermediate filament network perturbation in the *C. elegans* intestine causes systemic toxicity

**DOI:** 10.1101/2022.08.07.503108

**Authors:** Florian Geisler, Sanne Remmelzwaal, Mike Boxem, Rudolf E. Leube

**Affiliations:** Institute of Molecular and Cellular Anatomy, RWTH Aachen University, Aachen, Germany; Division of Developmental Biology, Institute of Biodynamics and Biocomplexity, Department of Biology, Faculty of Science, Utrecht University, Utrecht, The Netherlands

**Keywords:** Intermediate filament, endotube, intestine, *C*. *elegans*

## Abstract

Intermediate filaments (IFs) are major components of the metazoan cytoskeleton. A long-standing debate concerns the question whether IF network organization only reflects or also determines cell and tissue function. Using *C. elegans*, we have recently described mutants of the MAPK SMA-5, which perturb the organization of the intestinal IF cytoskeleton resulting in luminal widening and cytoplasmic invaginations. Besides these structural phenotypes, systemic dysfunctions were also observed. We now identify the IF polypeptide IFB-2 as a highly efficient suppressor of both the structural and functional deficiencies by removing the aberrant IF network. Mechanistically, IF network morphogenesis is linked to the phosphorylated IFB-2 aminoterminus. The rescuing capability is IF isotype-specific and not restricted to SMA-5 mutants but extends to other regulators of IF network morphogenesis, i.e. the cytoskeletal linker IFO-1 and the IF-associated protein BBLN1. The findings provide strong evidence for a gain-of-toxic function of the deranged IF networks with implications for diseases that are characterized by altered IF network organization.

## Introduction

Intermediate filaments (IFs) together with actin filaments and microtubules are important components of the cytoskeleton. They mediate mechanical tissue stability and have been implicated in multiple cellular processes such as vesicle transport, organelle positioning, cell cycle regulation, differentiation, metabolism, motility and stress response (Etienne-Manneville, 2018, Jacob et al., 2018, Leube and Schwarz, 2016, Margiotta and Bucci, 2016, Yoon and Leube, 2019, Coch et al., 2020, Toivola et al., 2010, Geisler and Leube, 2016). An ongoing debate is whether IFs are simply bystanders of these cellular processes or contribute actively to cell function and dysfunction (Etienne-Manneville, 2018, Bott and Winckler, 2020).

To study the morphogenesis and function of the IF system, we use the nematode *Caenorhabditis elegans*. A striking example of unique IF network organization is encountered in the intestine, where six IF polypeptides, i.e. IFB-2, IFC-1, IFC-2, IFD-1, IFD-2 and IFP-1, co-localize in the apical cytoplasm forming the electron dense endotube, which surrounds the lumen as a compact fibrous sheath and is attached to the composite *C. elegans* apical junction (CeAJ) (Carberry et al., 2009, Jahnel et al., 2016, Bossinger et al., 2004). It is assumed that this evolutionary conserved localization of the intestinal IF network (cf. (Coch and Leube, 2016)) mediates protection against mechanical stress (Geisler and Leube, 2016, Toivola et al., 2010, Geisler et al., 2016). Accordingly, the endotube is positioned at the interface between the cortical actin cytoskeleton with the stiff microvillar brush border and the soft cytoplasm (Geisler et al., 2020, McGhee, 2007, Bossinger et al., 2004). Because of its high degree of elasticity the IF-rich endotube likely dampens mechanical stresses occurring during food intake, defecation and body movement (Geisler et al., 2020). Elimination of the intestinal IFs IFB-2, IFC-2 and IFD-2 therefore leads to luminal widening although loss of IFC-1, IFD-1 and IFP-1 does not (Geisler et al., 2020). Ultrastructural analyses further showed that IFC-2 mutants have a rarefied endotube, whereas IFB-2 mutants lack it completely (Geisler et al., 2019, Geisler et al., 2020). The intestinal IF mutants present very mild organismal phenotypes with only minor or no detectable effects on development, progeny production, survival and stress sensitivity with the exception of IFC-2 mutants that are also expressed in the excretory canal and therefore induce more pronounced deficiencies (Geisler et al., 2020, Al-Hashimi et al., 2018, Carberry et al., 2009, Karabinos et al., 2001, Karabinos et al., 2004).

Modulators of intestinal IF distribution have been identified and characterized in *C. elegans* (Carberry et al., 2012, Geisler et al., 2016, Stutz et al., 2015, Estes et al., 2011, Geisler et al., 2019, Remmelzwaal et al., 2021, Koyuncu et al., 2021). One of them is SMA-5, a stress-activated kinase orthologue of the mitogen-activated protein kinase (MAPK) type (Watanabe et al., 2005). Abundant cytoplasmic invaginations of the adluminal, apical plasma membrane develop over time in *sma-5* mutant intestines (Geisler et al., 2016). These changes correlate with the development of a locally thickened endotube consisting of amorphous material next to areas with complete endotube loss. The cytoplasmic invaginations form at the transition between both areas. The structural changes go along with biochemical changes, i.e. altered IFB-2 phosphorylation. Furthermore, in comparison to the wild type and to intestinal IF mutants *sma-5* mutants are smaller, produce less offspring, develop more slowly, live shorter and are more sensitive to microbial pathogens and osmotic as well as oxidative stress (Geisler et al., 2016, Geisler et al., 2019). It is not known, whether these pathologies are attributable to the altered IF cytoskeleton or other *sma-5(n678)*-dependent cellular perturbations. Another modulator of the intestinal IF cytoskeleton is the intestinal intermediate filament organizer gene *ifo-1* (Carberry et al., 2012), which was originally identified as a cellular defense gene against pore-forming toxins and as part of the MAPK/JNK defense network (referred to as *ttm-4* in (Kao et al., 2011)). The IF network collapses in loss-of-function *ifo-1* mutants into large aggregates, which accumulate primarily at the CeAJ and occasionally in the cytoplasm. *ifo-1* mutants are small, have reduced progeny and are hypersensitive to different types of stress. The phenotypes are not only much more pronounced than those observed in IF mutants but are also more pronounced than in *sma-5* mutants (Geisler et al., 2019, Geisler et al., 2020, Carberry et al., 2012). Again, the contribution of the deranged IF network to the *ifo-1* mutant phenotype has not been determined to date. Recently, we described a third type of intestinal IF regulator, namely BBLN-1 (Remmelzwaal et al., 2021). BBLN-1 is a small coiled-coil protein whose depletion results in an intestinal phenotype that is highly reminiscent of that observed in *sma-5* mutants presenting bubble-shaped cytoplasmic invaginations of the apical plasma membrane. Systemic consequences of *bbln-1* mutation have not been analyzed in detail to date. Remarkably, all three regulators have been localized to the apical submembrane compartment in intestinal cells (Remmelzwaal et al., 2021, Geisler et al., 2016, Carberry et al., 2012).

To identify components of the IF-regulatory pathways in the *C. elegans* intestine, we performed a suppressor screen of *sma-5(n678)* mutants, which identified *ifb-2* mutation as the most efficient suppressor. Loss-of-function *ifb-2* mutation also partially rescued the *ifo-1* and *bbln-1* phenotypes. The perturbed intestinal IF networks were completely depleted in all instances. Even more importantly, the systemic dysfunctions were also rescued for the most part. These findings provide new evidence on the gain-of-toxic-function hypothesis driving the pathogenesis of aggregate-forming diseases, which have been shown to involve aberrant IF networks (e.g., (Coulombe et al., 2009, Chamcheu et al., 2011, Yoshida and Nakagawa, 2012, Clemen et al., 2013, Gentil et al., 2015, Didonna and Opal, 2019).

## Results

### The loss-of-function phenotype of the MAP-kinase SMA-5 is rescued by depletion of the intestinal intermediate filament polypeptide IFB-2

To identify downstream targets of the MAP-kinase orthologue SMA-5, we performed a genome-wide suppressor screen in OLB18 carrying the loss-of-function *sma-5(n678)* allele. Selecting for fast developing lines we isolated two lines, in which we identified an 83 bp deletion in the *ifb-2* gene (positions 5.751.812-5.751.893; allele *ifb-2(kc20)*). The mutant *ifb-2* gene encodes 120 amino acids, 116 of which correspond to the aminoterminal amino acids of IFB-2. The fragment encompasses the head domain together with 73 amino acids of the coiled-coil rod domains 1a and 1b (Karabinos et al., 2004). To demonstrate that loss-of-function of IFB-2 is by itself sufficient and necessary to rescue the *sma-5(n678)* phenotype and to exclude that the residual aminoterminal fragment that is still produced from *kc20* is responsible for the rescue, we crossed *sma-5(n678)* mutants with the previously described *ifb-2* knockout allele *kc14* (Geisler et al., 2019). The *kc14* allele encodes a 29 amino acid-long oligopeptide encompassing only 13 of the most aminoterminal amino acids of IFB-2. Assessment of body length revealed a near normal body size of the double *sma-5(n678);ifb-2(kc14)* mutants. Only a minor reduction was detected at day 4 as was the case for *ifb-2(kc14)* single mutants (Fig. S1). Normal body length was reached by day 5 in *sma-5(n678);ifb-2(kc14)* and *ifb-2(kc14)* but not in *sma-5(n678)* (not shown).

Light microscopy revealed a rescue of the luminal widening and cytoplasmic invagination phenotypes (Fig. 1 A-D). Ultrastructural analyses further showed a reversion of the phenotype to a near wild-type situation in the double mutants (Fig. 1 E-H). Only minor luminal undulations and slight perturbation of the microvillar brush border remained. Most notably, the endotube was completely absent (Fig. 1 H). The ultrastructural features were identical to those encountered in *ifb-2(kc14)* single mutants (Fig. 1 F).

**Figure 1.**
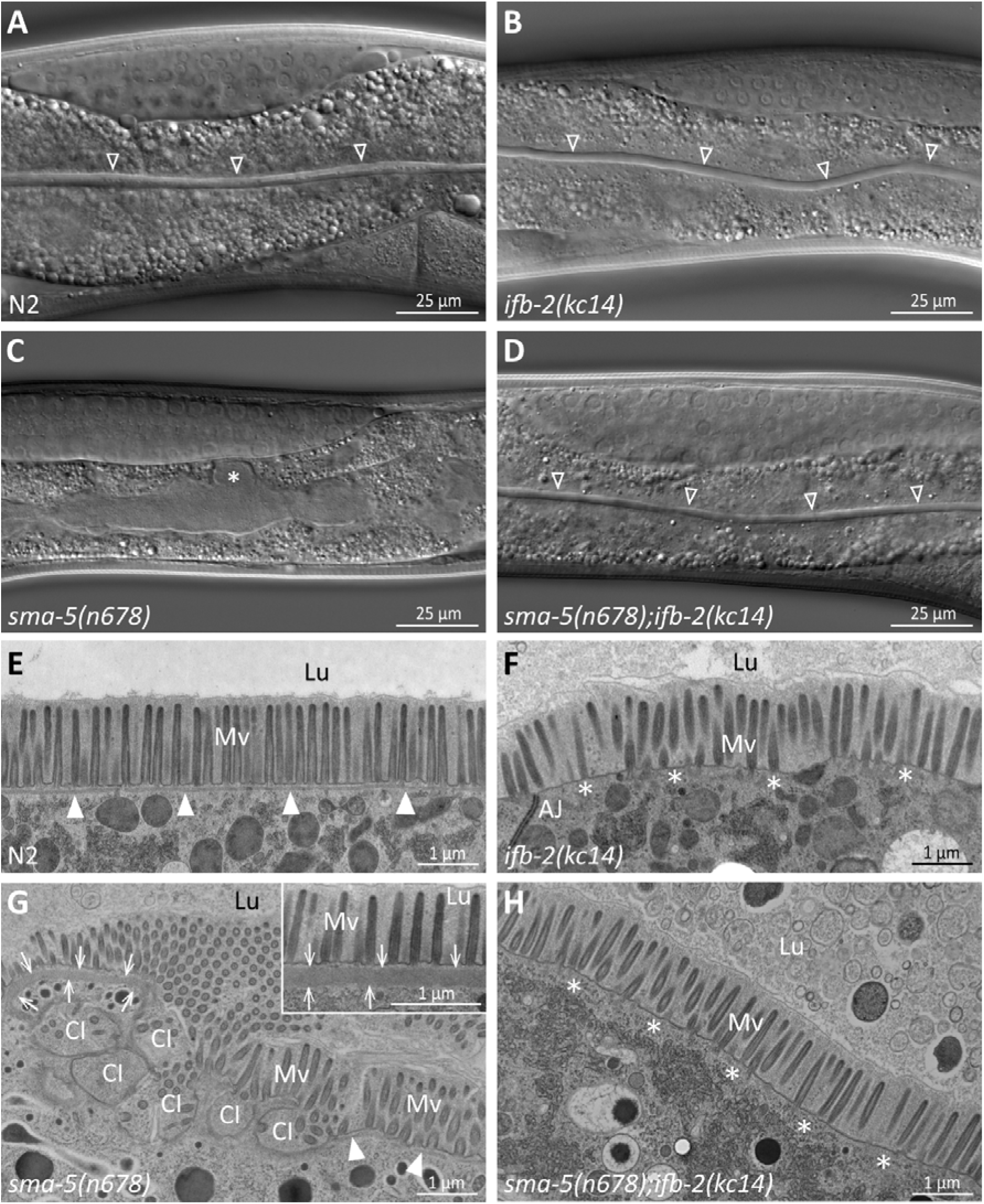
The *ifb-2(kc14)* knockout allele rescues the luminal widening and cytoplasmic invagination phenotypes and abolishes the aberrant IF network of *sma-5(n678)* mutants. **(A-D)** Differential interference contrast pictures of viable wild-type N2 **(A)**, *ifb-2(kc14)* **(B)**, *sma-5(n678)* **(C)** and *sma-5(n678);ifb-2(kc14)* animals **(D)**. *ifb-2* knockout causes only minor intestinal defects. In contrast, *sma-5(n678)* animals display extensive luminal widening and large cytoplasmic invaginations of the apical plasma membrane in intestinal cells (asterisk in **C**). This phenotype is effectively reversed by the *ifb-2(kc14)* knockout allele. Non-filled arrowheads: normal-appearing intestinal lumen. **(E-H)** Electron microscopy images of high-pressure frozen samples show wild-type N2 **(E)**, *ifb-2(kc14)* **(F)**, *sma-5(n678)* **(G)** and *sma-5(n678);ifb-2(kc14)* intestinal cell apices **(H)**. *sma-5(n678)* animals contain regions with an enlarged endotube consisting of densely packed IFs (arrows) and differently-sized cytoplasmic invaginations (CI) with no endotube or a reduced endotube (arrowheads). Additional knockout of *ifb-2* almost restores the wild-type morphology with only residual luminal widening and mildly perturbed microvillar arrangement. But the endotube, which is easily detected in the wild type (filled arrowheads), is completely absent in single *ifb-2(kc14)* and double *sma-5(n678);ifb-2(kc14)* mutants (expected position marked by asterisks). Lu, lumen; Mv, microvilli; AJ, *C. elegans* apical junction.

The *ifb-2(kc14)* knockout allele furthermore restored the prolonged time development of *sma-5(n678)* (Fig. 2 A). It was, however, still elevated in comparison to the wild type and was only slightly different from the single *ifb-2(kc14)* mutant. To define minor alterations in development more precisely, analysis of the different developmental stages was carried out next. The results shown in Fig. 2 B illustrate the high degree of similarity in the developmental time course of the single *ifb-2(kc14)* and double *sma-5(n678);ifb-2(kc14)* mutants. Both developed more slowly than the wild type and much faster than the *sma-5(n678)* animals, some of which never reached adulthood. The latter could be ascribed to larval arrest at the L4 stage (Fig. 2 C).

**Figure 2.**
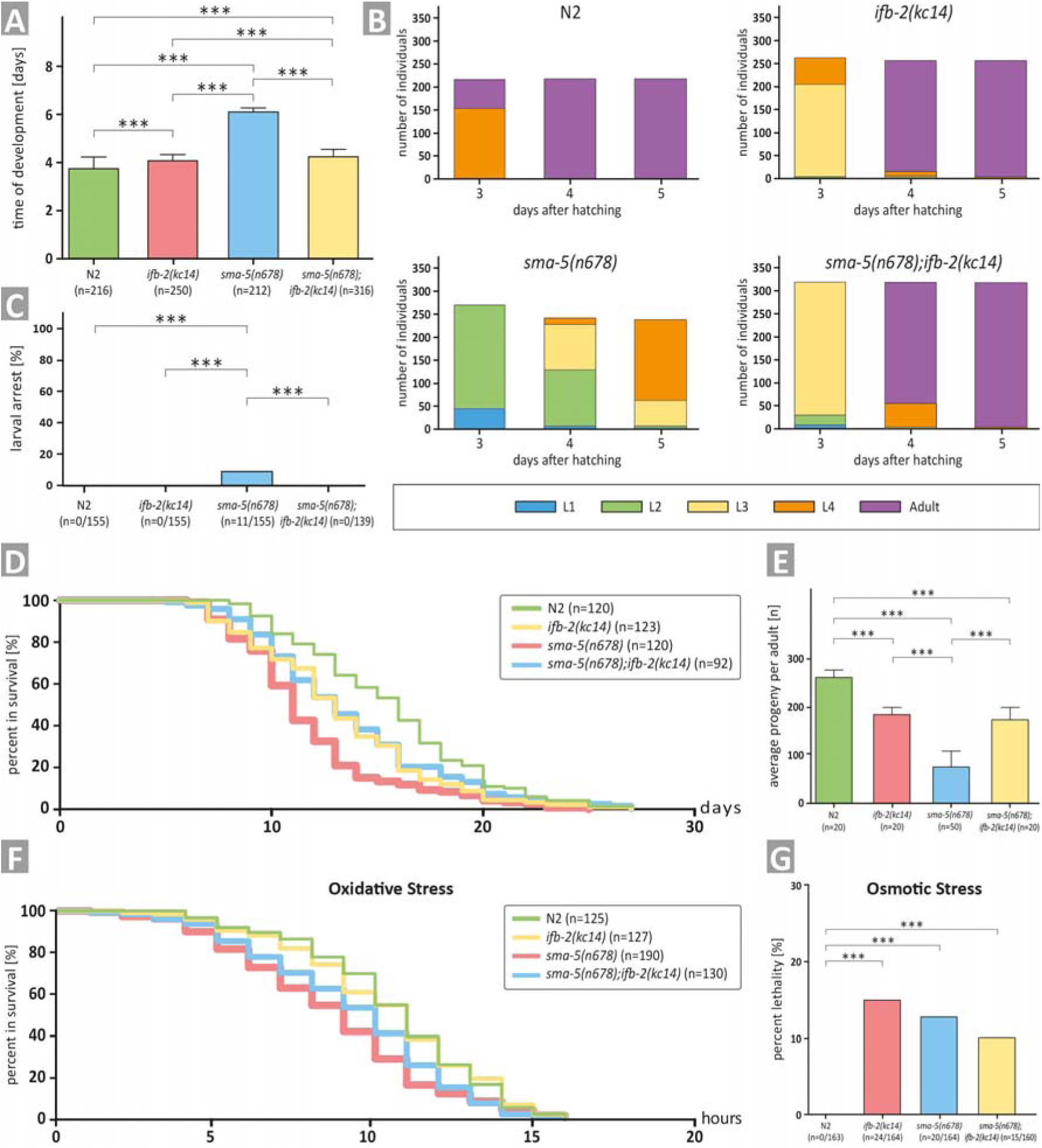
Depletion of IFB-2 (*ifb-2(kc14)*) rescues developmental retardation, larval arrest, reduced median life span, decreased brood size and increased sensitivity to oxidative stress of *sma-5(n678)* mutants. **(A)** The histogram shows a comparison of the time of development in N2, *ifb-2(kc14)*, *sma-5(n678)* and *sma-5(n678);ifb-2(kc14)* (N2: 3.7±0.5 days; *ifb-2(kc14)*: 4.1±0.2 days; *sma-5(n678)*: 6.0±0.2 days; *sma-5(n678);ifb-2(kc14)*: 4.2±0.4 days; *** p<0.0001). **(B)** The color-coded histogram depicts the number of larval and adult stages detected 3, 4 and 5 days after hatching. **(C)** The histogram illustrates a complete rescue of the larval arrest phenotype observed in *sma-5(n678)* by *ifb-2(kc14)* (N2: 0%; *ifb-2(kc14)*: 0%; *sma-5(n678)*: 7.1%; *sma-5(n678);ifb-2(kc14)*: 0%). **(D)** The plot shows that the reduced life span of *sma-5(n678)* is rescued by addition of *ifb-2(kc14)* to the level encountered in *ifb-2(kc14)* but not the wild-type level (median survival for N2: 16 days; *ifb-2(kc14)*: 13 days; *sma-5(n678)*: 11 days; *sma-5(n678);ifb-2(kc14)*: 13 days; p=0.0004 *ifb-2(kc14)* versus N2; p<0.0001 *sma-5(n678)* versus N2; p=0.0003 *sma-5(n678)* versus *ifb-2(kc14)*; p<0.0001 *sma-5(n678);ifb-2(kc14)* versus N2; p<0.0046 *sma-5(n678);ifb-2(kc14)* versus *sma-5(n678)*). **(E)** The histogram reveals that the drastic reduction in progeny observed in *sma-5(n678)* mutants versus N2 (76±31 versus 263±13; p<0.0001) is rescued in *sma-5(n678);ifb-2(kc14)* double mutants (175±21 versus 76±31; p<0.001) but does not reach wild-type level (175±21 versus 263±13; p<0.001) and is similar to *ifb-2(kc14)* (175±21 versus 183±15; p>0.5). **(F)** The survival plot shows the effect of acute oxidative stress in the wild type (N2), *ifb-2(kc14)*, *sma-5(n678)* and *sma-5(n678);ifb-2(kc14)* backgrounds (median survival for N2: 11 h; *ifb-2(kc14)*: 11 h; *sma-5(n678)*: 9 h; *sma-5(n678);ifb-2(kc14)*: 10 h; p<0.0001 for N2 or *ifb-2(kc14)* versus *sma-5(n678)*; p<0.05 for *sma-5(n678);ifb-2(kc14)* versus *sma-5(n678)*; p<0.01 for N2 or *ifb-2(kc14)* versus *sma-5(n678);ifb-2(kc14)*). **(G)** The histogram scores the percentage of dead worms in response to acute osmotic stress for N2 (0%), *ifb-2(kc14)* (14.6%), *sma-5(n678)* (12.2%) and *sma-5(n678);ifb-2(kc14)* (9.4%).

Life span determinations further showed that the shortened median life span of *sma-5(n678)* animals could be rescued in the *sma-5(n678);ifb-2(kc14)* double mutants to the level observed in *ifb-2(kc14)* single mutants. Yet, the life spans of *sma-5(n678);ifb-2(kc14)* and *ifb-2(kc14)* mutants were still reduced in comparison to the wild type by one day (Fig. 2 D). Similarly, the drastic reduction in progeny of *sma-5(n678)* was rescued in the double mutant to the level observed in *ifb-2(kc14)*, which, again, was significantly less than in the wild type (Fig. 2 E).

In a next set of experiments, we investigated whether loss of IFB-2 would also rescue the increased stress sensitivity of *sma-5(n678)* (Geisler et al., 2019). In the presence of oxidative stress, the survival *of sma-5(n678)* was reduced by 3.5 h in comparison to wild-type N2 but was only reduced by 0.5 h and 1.5 h in *ifb-2(kc14)* and *sma-5(n678);ifb-2(kc14)*, respectively (Fig. 2 F). All mutants appeared to be similarly affected in osmotic stress assays and a statistically significant rescue phenotype could not be observed (Fig. 2 G).

Taken together, we can conclude that loss of IFB-2 rescues all major SMA-5 phenotypes to levels observed in *ifb-2(kc14)*. This demonstrates that the alterations of the IF network observed in *sma-5(n678)* exert a gain-of-toxic function by negatively affecting intestinal and organismal physiology.

### Depletion of individual intestinal intermediate filament polypeptides reveals isotype-specific rescue efficiency of the *sma-5* phenotype

To test whether the observed rescue is specific for IFB-2 or applies also to the other five IFs that are expressed in the intestine, each IF was downregulated by RNAi in the *sma-5(n678)* background. As simple readouts, F1 progeny from RNAi-treated worms was imaged on agar plates four days after egg-laying (Fig. 3 A-G) and the body length was measured (Fig. 3 H). The assays confirmed the expected efficient rescue of the developmental growth defect in *sma-5(n678)* by *ifb-2(RNAi)*. An easily detectable, though reduced rescuing efficiency was detected for *ifc-2(RNAi)*. An even less pronounced but statistically still significant rescue could be identified in *ifd-1*, *ifd-2* and *ifp-1* RNAi-treated animals and none for *ifc-1(RNAi)*.

**Figure 3.**
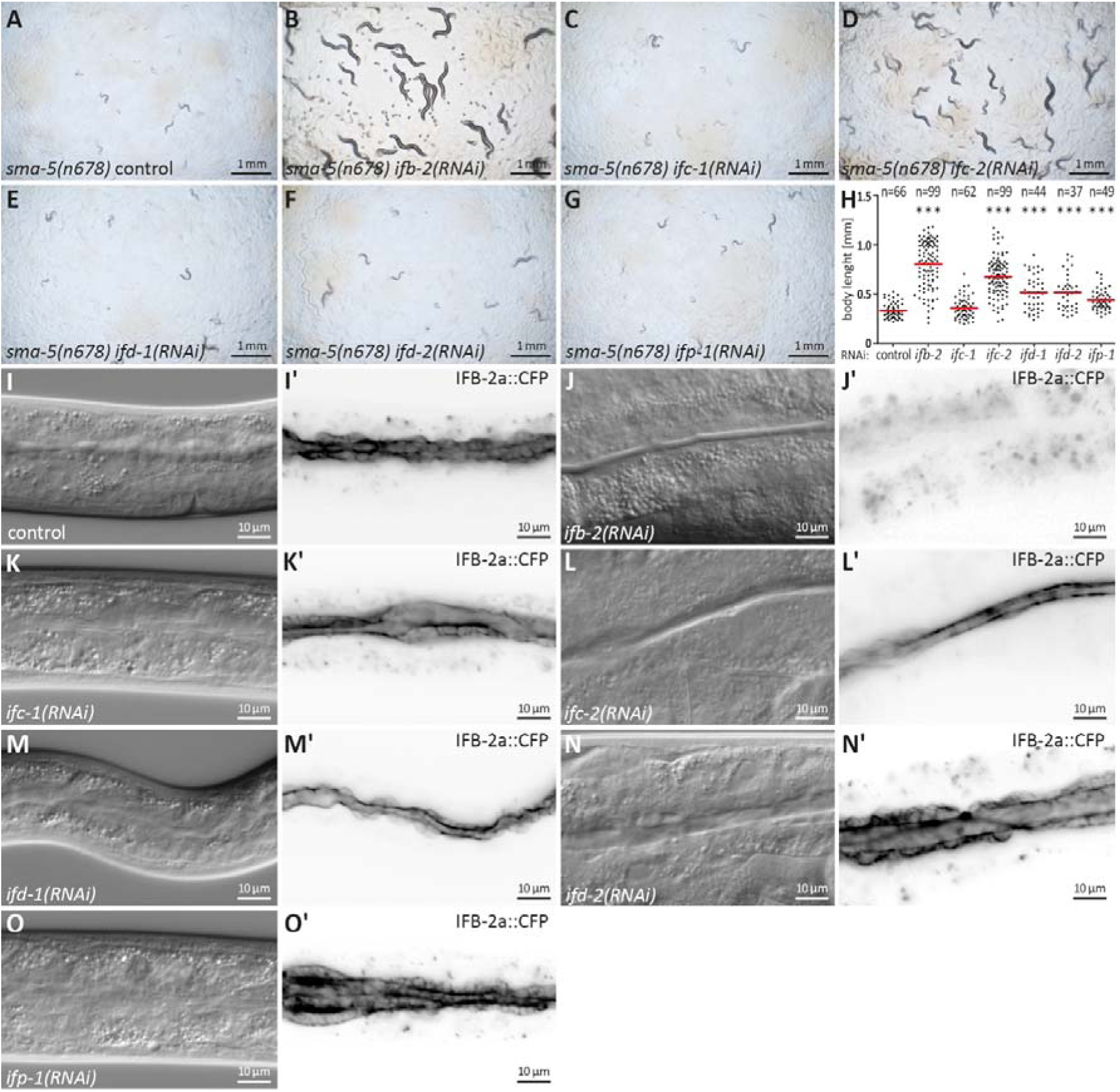
Downregulation of *ifb-2* and *ifc-2* is most efficient in suppressing developmental retardation, small body size and cytoplasmic invaginations of *sma-5(n678)*. **(A-G)** show bright field images of agar plates containing F1 *sma-5(n678)* subjected to RNAi 4 days after egg-laying as indicated. **(H)** The scatter dot blot summarizes the results of body length measurements in *sma-5(n678)* subjected to either empty RNAi vector (control; 331±72.71 µm) or *ifb-2(RNAi)* (802.90±235.10 µm), *ifc-1(RNAi)* (358.50±103.50 µm), *ifc-2(RNAi)* (675.60±194.10 µm), *ifd-1(RNAi)* (511.70±172.90 µm), *ifd-2(RNAi)* (512.60±178.80 µm) or *ifp-1(RNAi)* (443.30±109 µm). Note that the strongest rescue is observed for *ifb-2* followed by *ifc-2* and trailed by *ifd-1*, *ifd-2* and *ifp-1* all of which are statistically significant (p<0.0001). No detectable rescue is observed for *ifc-1*. **(I-O’)** The microscopic images show differential interference contrast at left and corresponding fluorescence detection (inverse presentation) of the IFB-2a::CFP reporter in vital *sma-5(n678)* animals after RNAi against *ifb-2* **(J, J’)**, *ifc-1* **(K, K’)**, *ifc-2* **(L, L’)**, *ifd-1* **(M, M’)**, *ifd-2* **(N, N’)** and *ifp-1* **(O, O’)** (RNAi control in **I, I’**). Note that the knockdown of *ifb-2* leads to a loss of reporter fluorescence and efficiently rescues the *sma-5(n678)* invagination phenotype. A rescue is also detectable after loss of IFC-2 albeit at reduced efficiency. None of the other knockdowns resulted in detectable reduction of the invagination phenotype.

We next examined whether the cytoplasmic invagination phenotype could be rescued in a similar fashion. To this end, the different IF-encoding RNAs were downregulated in the IFB-2a::CFP reporter strain (Fig. 3 I-O’). As predicted, *ifb-2(RNAi)* abolished the reporter fluorescence as well as the cytoplasmic invaginations of *sma-5(n678)*. *ifc-2(RNAi)* also led to a reduction of the cytoplasmic invagination phenotype but was much less efficient than *ifb-2(RNAi)*. All other interfering RNAs, however, did not visibly affect the invagination phenotype. Nevertheless we cannot exclude that the presence of the reporter obscured the effect of RNA downregulation of these IF polypeptides.

Taken together, the isotype-specific phenotypes may be explained by the abundance and polymerization of the different IF polypeptides. Thus, IFB-2 has been shown to pair with multiple intestinal IFs (Karabinos et al., 2017) and has been identified as the master regulator of the endotube (Geisler et al., 2020). It is therefore not surprising that IFB-2 depletion elicits the strongest rescuing activity of the *sma-5(n678)* mutant phenotype. It is also in line with IFC-2 depletion being second in line as also evidenced in previously reported effects on endotube structure and stress sensitivity (Geisler et al., 2020). Overall, our findings demonstrate that the altered IF network and not a single IF is responsible for the observed rescue of the *sma-5(n678)* phenotype.

### The phosphorylated aminoterminal domain of IFB-2 contributes to intestinal IF network morphogenesis and function

In search for a molecular mechanism that may be involved in altered IF network morphogenesis, it is of interest that the head domains of mammalian IF polypeptides have been implicated in cytoplasmic IF assembly (Omary et al., 2006, Zhou et al., 2021, Hatzfeld and Burba, 1994). Given the essential contribution of IFB-2 to intestinal IF network formation, we concentrated on this polypeptide. We deleted base pairs 4-1107 of the *ifb-2* gene by CRISPR-cas resulting in a truncated protein missing the complete aminoterminal head domain (*ifb-2(mib169[S2-M43 deleted])II*) (cf. (Karabinos et al., 2004)). Mutants showed a phenotype reminiscent of that described for *sma-5(n678)* animals with cytoplasmic invaginations of the intestinal lumen accompanied by a rarefied IFC-2 fluorescence with regions completely lacking a signal next to areas showing enhanced fluorescence (Fig. 4 A-B’). These observations provide evidence for the evolutionary conserved importance of the IF head domain for IF network formation.

**Figure 4.**
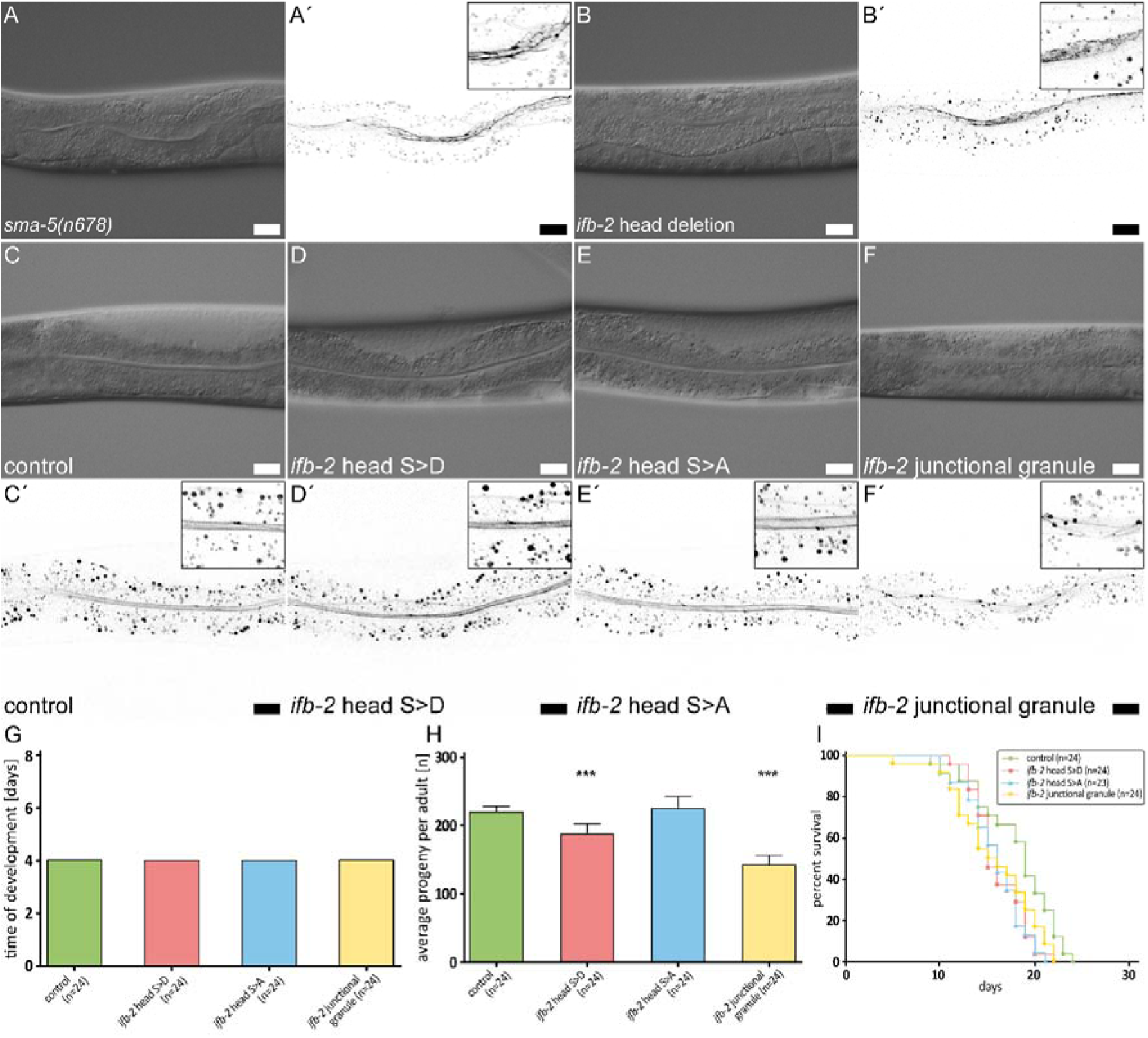
The phosphorylated IFB-2 aminoterminus is involved in intestinal IF network morphogenesis, progeny production and life span. **(A-F’)** The microscopy images show differential interference contrasts **(A-F)** and corresponding fluorescence recordings of the IFC-2a/e::YFP reporter (inverse presentation; **A’-F’**) in vital *sma-5(n678)* **(A-A’)**, *ifb-2* head deletion **(B-B’**, *ifb-2(mib169[D2-M43 deleted])II*), control **(C-C’)**, phosphomimetic *ifb-2* head S>D **(D-D’**, *ifb-2(kc22[S2D;S5D;S7D;S16-19D;S24D;S35D])II*), phosphodeficient *ifb-2* head S>A **(E-E’**, *ifb-2(kc26[S2A;S5A;S7A;S16-19A;S24A;S35A])II*) and junctional granule mutants **(F-F’**, *ifb-2(kc27[S2A;S5A;S7A;S16-19A;E31-A184 deleted])II*). Scale bars: 20 µm. **(G-H)** The histograms present a comparison of time of development **(G)** and average progeny **(H)** of control, phosphomimetic *ifb-2* head S>D, phosphodeficient *ifb-2* head S>A and junctional granule mutants. None of the mutants shows a prolonged development (control: 4.0 days; *ifb-2* head S>D: 4.0 days; *ifb-2* head S>A: 4.0 days; junctional granule mutant: 4.0 days). The average progeny per adult, however, is reduced in *ifb-2* head S>D but not in *ifb-2* head S>A mutants in comparison to control (control: 227±9; *ifb-2* head S>D: 193±16; *ifb-2* head S>A: 232±18; p<0.0001 control versus *ifb-2* head S>D). Note the even higher reduction in progeny observed in junctional granule mutants versus control (control: 227±9; junctional granule mutant: 145±16; p<0.0001). **(I)** The survival plot shows that mutants with genetic modifications of the *ifb-2* head domain have a reduced life expectancy albeit to different degrees with the *ifb-2* head S>D mutant showing the most severe phenotype (median survival for control: 19 days; *ifb-2* head S>D: 15 days; *ifb-2* head S>A: 16 days; junctional granule mutant: 15.5 days; p=0.0471 *ifb-2* head S>D versus control; p=0.0206 *ifb-2* head S>A versus control; p=0.0468 junctional granule mutant versus control; p<0.0001).

Previous observations had also shown that phosphorylation sites in the head domain of IFs are most frequently targeted by kinases with major consequences on IF network dynamics and organization (review in (Sawant and Leube, 2017, Snider and Omary, 2014)). In accordance, Wormbase and Phosida list multiple serine phosphorylation sites in the IFB-2 head domain. We therefore decided to mutate putative phosphorylation targets in the IFB-2 head domain by introducing phosphomimetic S>D and phosphorylation-deficient S>A mutations into all serines of the head region, namely S2, S5, S7, S16-S19, S24 and S35 (alleles *kc22* and *kc26*, respectively). Generating the mutant *ifb-2* alleles in the IFC-2a/e::YFP background allowed direct examination of the effect on intestinal IF network formation. Neither the phosphomimetic nor the phosphorylation-deficient mutants presented a visible change in IFC-2 localization (Fig. 4 C-E’). Further characterization revealed that the mutants showed no developmental delay (Fig. 4 G) and only a slight reduction of progeny in case of the *ifb-2* head S>D mutants (Fig. 4 H). However, significant reduction of life span could be observed in all instances (Fig. 4 I). Taken together, while the phosphorylation sites in the head domain of IFB-2 are not necessary for IF network morphogenesis *per se*, they may affect network properties such as subunit turnover that are not readily apparent from static pictures and may result in reduced network resilience leading to the decreased life span.

During screening of the different CRISPR mutants we noted an aberrant IFC-2 fluorescence pattern presenting a reduced though still apically localized IFC-2 distribution with additional, strongly fluorescent granules next to the CeAJ (referred to as junctional granule mutant, *ifb-2(kc27)*) (Fig. 4 F-F’). This distribution pattern is somewhat reminiscent of the *ifo-1* loss-of-function phenotype that is, however, characterized by much larger junctional aggregates (Carberry et al., 2012). Sequence analysis identified an E31-A184 deletion in combination with S2/S5/S7/S16-19>A mutations in IFB-2. The mutant protein could be detected by immunofluorescence in the intestine, albeit at highly reduced levels in the cytoplasm, the CeAJ and CeAJ-associated apical granules (Fig. S2 A-B’). The truncated IFB-2a and IFB-2b proteins of about 50 kDa were identified in immunoblots (Fig. S2 C). Significant effects on progeny and life expectancy were also noted (Fig. 4 H-I).

We conclude that the head domain together with the aminoterminal rod domain of IFB-2 are involved in phosphorylation-dependent IF network formation and function in the intestinal epithelium.

### *ifb-2(kc14)* partially rescues the *ifo-1(kc2)* and *bbln-1(mib70)* phenotypes

The above findings suggested that the rescue function of IFB-2 deletion in *sma-5(n678)* was caused by removal of the densely packed IFs. To test the hypothesis that perturbed IF network organization *per se* exerts a toxic effect, we studied another paradigm, namely the *ifo-1(kc2)* phenotype, which is characterized by prominent IF polypeptide-containing junctional aggregates (Carberry et al., 2012). To this end, we crossed *ifo-1(kc2)* with *ifb-2(kc14)*. As predicted, the double mutant lacked the junctional IF aggregates altogether (Figure 5 A-D). Occasionally, a faint but largely reduced remnant endotube structure was detectable, often in the vicinity of the CeAJ. Furthermore, double mutants had an almost normal intestinal lumen and inconspicuous microvilli (Fig. 5 D). Additional analyses showed a significantly reduced time of development of *ifo-1(kc2);ifb-2(kc14)* in comparison to *ifo-1(kc2)*, which was, however, significantly retarded in comparison to *ifb-2(kc14)* or wild-type N2 (Figure 5 E-I). Moreover, analysis of body-length revealed a significant rescue of the *ifo-1(kc2)* growth defect in the absence of the IFB-2 protein (Fig. S1).

**Figure 5.**
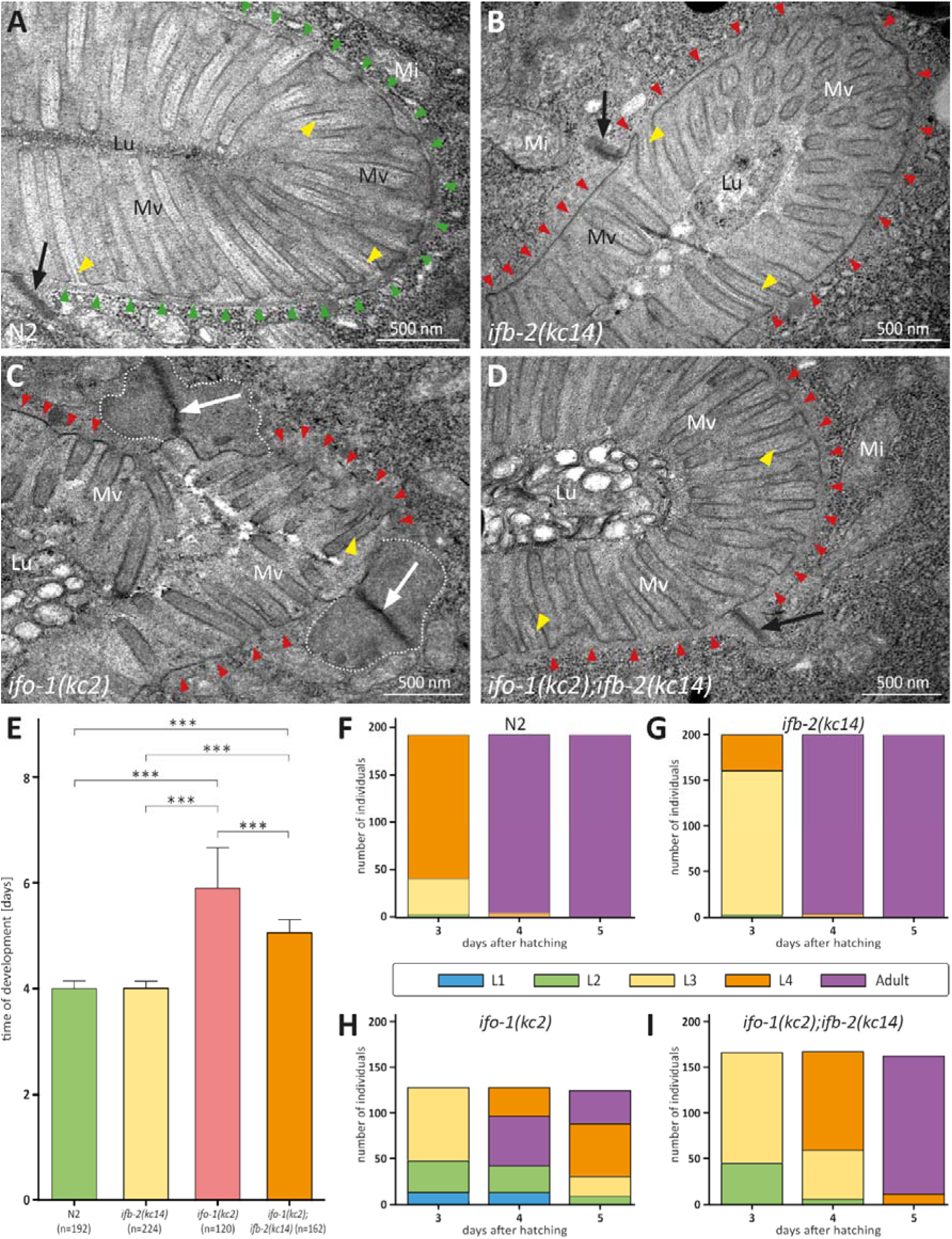
Depletion of IFB-2 partially rescues the *ifo-1(kc2)* phenotype. **(A-D)** The electron micrographs show a comparison of the intestinal cell apices surrounding the lumen (Lu) of wild-type N2 (A), *ifb-2(kc14)* (B), *ifo-1(kc2)* (C) and *ifo-1(kc2);ifb-2(kc14)* (D). Note the distinct endotube in N2 (green arrowheads) and its absence in *ifb-2(kc14)*, *ifo-1(kc2)* and *ifo-1(kc2);ifb-2(kc14)* (red arrowheads). The pathognomonic large junctional aggregates of *ifo-1(kc2)* are delineated by broken white lines. Note also the improved brush border morphology in D compared to C (Mv, microvilli). Arrows, CeAJ; yellow arrowheads, microvillar actin bundles; Mi, mitochondrion. **(E-I)** The histograms depict the time of development in E and the number of staged worms at different times after hatching in N2 (F), *ifb-2(kc14)* (G), *ifo-1(kc2)* (H) and *ifo-1(kc2);ifb-2(kc14)* (I). Note the partial rescue in the double mutants. (N2: 4.0±0.1 days; *ifb-2(kc14)*: 4.0±0.1 days; *ifo-1(kc2)*: 5.9±0.8 days; *ifo-1(kc2);ifb-2(kc14)*: 5.1±0.3 days; *** p<0.0001).

In a next set of experiments we analyzed the rescuing efficiency of IFB-2 depletion in *bbln-1* mutants, which present a structural phenotype similar to but not identical to that encountered in *sma-5* mutants (Remmelzwaal et al., 2021). It had been shown recently that depletion of IFB-2 rescues the intestinal cytoplasmic invagination phenotype of *bbln-1* mutants (Remmelzwaal et al., 2021). We now tested, whether systemic dysfunctions also occur in *bbln-1* mutants and whether they can be rescued by IFB-2 depletion. We found that the *bbln-1(mib70)* loss-of-function mutants exhibited systemic dysfunctions as determined by body size and time of development (Fig. S1, Fig. 6 A). Although BBLN-1 expression is, in contrast to IFO-1 and SMA-5, not limited to the intestine, the systemic dysfunctions were less pronounced than in *sma-5(n578)* or *ifo-1(kc2)*. The *ifb-2(kc14)* knockout allele rescued the reduced body size and prolonged time of development of *bbln-1(mib70)* (Fig. S1, Fig. 6 A). The remaining growth defect and developmental retardation, however, were still slightly increased not only in comparison to the wild type but also to single *ifb-2(kc14)* mutants. To define the alterations in development more precisely, we analyzed the different developmental stages every day after hatching. The results shown in Fig. 6 B-E highlight the high degree of similarity in the developmental time course of the single *ifb-2(kc14)* and double *bbln-1(mib70);ifb-2(kc14)* mutants. Both develop more slowly than the wild type and much faster than *bbln-1(mib70)* animals.

**Figure 6.**
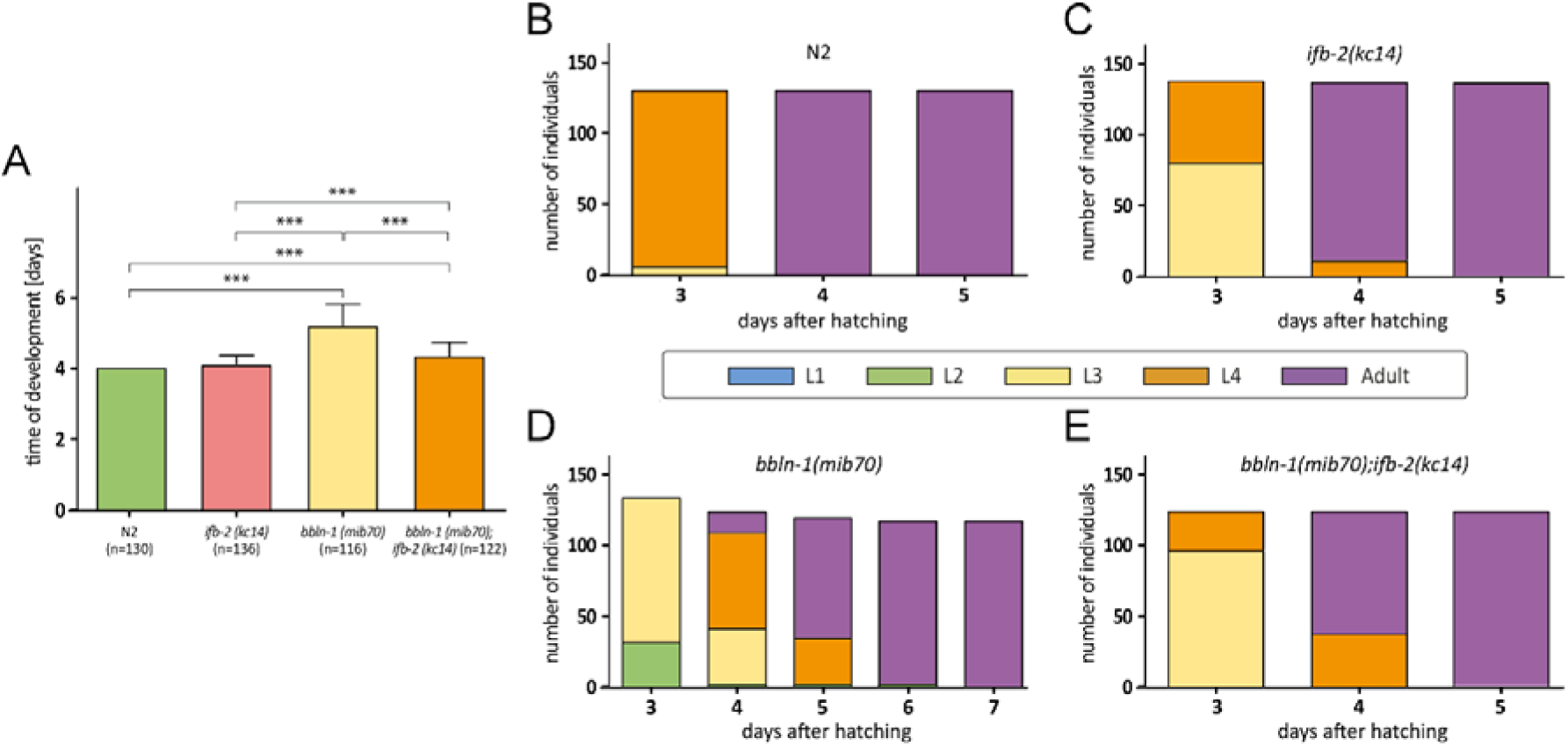
Depletion of IFB-2 partially rescues the *bbln-1(mib70)* phenotype. **(A-E)** The histograms depict the time of development in **(A)** and the number of staged worms at different times after hatching in N2 **(B)**, *ifb-2(kc14)* **(C)**, *bbln-1(mib70)* **(D)** and *bbln-1(mib70);ifb-2(kc14)* **(E)**. Note the partial rescue in the double mutants. (N2: 4.0 days; *ifb-2(kc14)*: 4.1±0.3 days; *bbln-1(mib70)*: 5.2±0.6 days; *bbln-1(mib70);ifb-2(kc14)*: 4.3±0.5 days; *** p<0.0001).

Fig. 7 summarizes the morphological observations in the intestine for the three types of IF regulators together with the rescue phenotype resembling that of the *ifb-2(kc14)* mutant. It draws attention to the major conclusion of the manuscript that perturbed IF networks affect the structural and functional integrity of the intestinal network, the removal of which alleviates the mutant phenotypes.

**Figure 7.**
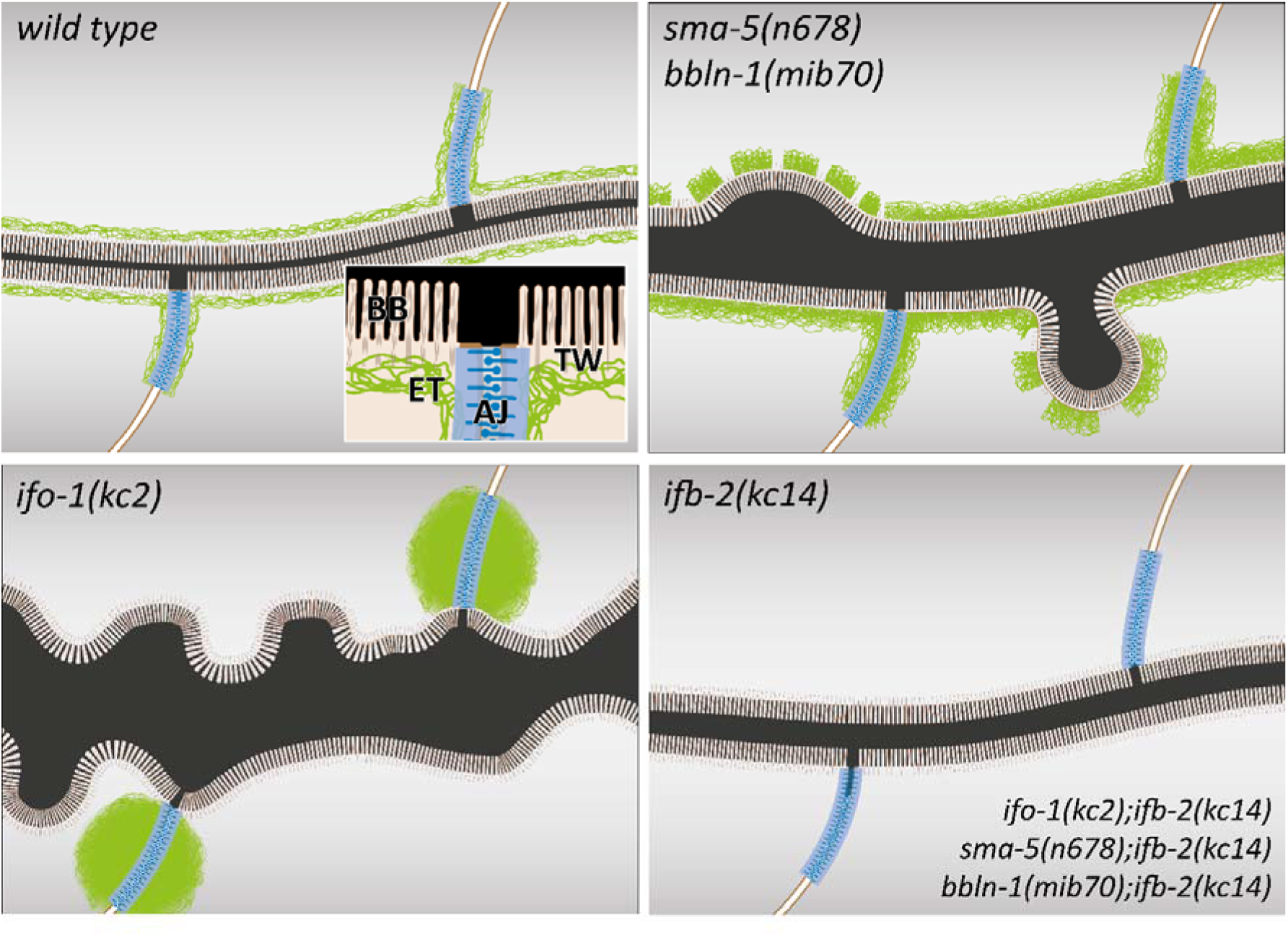
The schematic drawings highlight changes in the intestinal lumen and IF network organization in the different mutant backgrounds. The wild-type scheme (upper left) depicts the adluminal brush border (BB) consisting of microvilli with bundled actin filaments, the terminal web (TW) with traversing microvillar actin rootlets that rest on the IF-rich endotube (ET) together with the *C. elegans* apical junction (AJ), which serves as an anchorage site for the IF network. Note the luminal enlargement and cytoplasmic invaginations in *sma-5(n678)* and *bbln-1(mib70)* presenting a thickened and discontinuous endotube with slightly disordered microvilli (upper right). *ifo-1(kc2)* is characterized by endotube loss and formation of intermediate filament aggregates at the *C. elegans* apical junction (lower left). Luminal widening, cytoplasmic invaginations and microvillar disorder are also observed. The mildest phenotype is detectable in *ifb-2(kc14)* and in the double mutants *ifo-1(kc2)*;*ifb-2(kc14)*, *sma-5(n678)*;*ifb-2(kc14)* and *bbln-1(mib70)*;*ifb-2(kc14)*.

## Discussion

Exploiting rapid *C. elegans* genetic screening, we observed that the complex phenotypes induced by mutation of the MAPK orthologue SMA-5 can be rescued by deletion of the cytoskeletal IF protein IFB-2. It included rescue of structural defects (cytoplasmic invagination and lumen dilatation), developmental and growth defects as well as oxidative stress resilience. The obtained rescue levels coincided precisely with the mild phenotype of single *ifb-2(kc14)* mutants. Remarkably, osmotic stress resilience could not be rescued suggesting that this loss-of-function is tightly coupled to the absence of the endotube. In accordance, it has been suggested that osmotic stress resilience is a fundamental function of cytoplasmic IFs (D’Alessandro et al., 2002, Pekny and Lane, 2007). Our findings furthermore provide strong *in vivo* evidence that the rescued *sma-5* mutant phenotype is caused by the presence of IFB-2-containing pathological assemblies. This conclusion was confirmed in *ifo-1* and *bbln-1* mutants. Removal of the pathological IFB-2 assemblies also rescued complex biological functions. We therefore conclude that the deranged intestinal IF cytoskeletons are associated with a gain-of-toxic function exerting toxic effects, which have detrimental consequences for cell and tissue function and thereby adversely affect growth and reproduction of the entire organism.

The fact that removal of the very differently deranged IFs, i.e. the thick subapical slabs in *sma-5(n678)*) and *bbln-1(mib70)* and the large junctional aggregates in *ifo-1(kc2)*, rescued the phenotypes in all instances is quite remarkable since the mechanical and structural dysfunctions appear to be fundamentally different in the different mutant backgrounds. They manifest in *sma-5* and *bbln-1* mutants as prominent cytoplasmic invaginations and primarily as luminal widening in *ifo-1* mutants (see also (Geisler et al., 2016, Carberry et al., 2012, Remmelzwaal et al., 2021)). Furthermore, the absence of an endotube in *ifo-1* mutants predicts that the force equilibrium is affected differently from that in *sma-5* and *bbln-1* mutants with the locally thickened endotube. This indicates that restored mechanics alone are not sufficient to explain the phenotypic restoration in the double mutants. Thus, other non-mechanical functions must be attributable to the abnormal IF assemblies. It is interesting in this context that IFs may serve as signaling platforms by providing a large scaffold capable of sequestering and positioning signaling molecules that can be recruited by weak interactions and can be released, e.g., by protein modification or structural changes of the IF cytoskeleton (review in Bott and Winckler, 2020, Coulombe and Omary, 2002, Magin et al., 2007). Future experiments will show, whether such a scaffolding function is compromised in *sma-5*, *bbln-1* and *ifo-1* mutants by either sequestering or setting free regulatory factors that modulate pathways needed for normal growth, development and stress responses.

Our findings are likely relevant for the multiple aggregate-forming human diseases that involve IF polypeptides (e.g., (Coulombe et al., 2009, Chamcheu et al., 2011, Yoshida and Nakagawa, 2012, Clemen et al., 2013, Gentil et al., 2015, Didonna and Opal, 2019). They mandate a careful evaluation of possible toxic functions of the perturbed IF networks, which may not be limited to the local cell and tissue environment but may have systemic consequences.

## Material and Methods

### *C. elegans* strains and bacteria

Wild-type strain N2, strain FK312 *sma-5(n678)X* and OP50 bacteria were obtained from the Caenorhabditis Genetics Center (CGC; University of Minnesota, MN, USA). Strains BJ49 *kcIs6[ifb-2p::ifb-2a::cfp]IV* (Hüsken et al., 2008), BJ142 *ifo-1(kc2)IV* (Carberry et al., 2012), OLB18 *sma-5(n678)X;kcIs6[ifb-2p::ifb-2a::cfp]IV* (Geisler et al., 2016), BJ309 *ifb-2(kc14)II* (Geisler et al., 2019) and BOX415 *erm-1(mib40[erm-1::AID::mCherry])I;Pelt-2::TIR::tagBFP-2::tbb-2-3’UTR (mib58[Pelt-2::TIR-1::tagBFP-2-Lox511::tbb-2-3’UTR])IV;C15C7.5(mib70[pC15C7.5::GFP1-3, C15C7.5 0-914del])X* (Remmelzwaal et al., 2021) have been described. Strain FK312 was crossed with strain BJ309 to obtain strain BJ346 *sma-5(n678)X;ifb-2(kc14)II*. Strain BJ328 *ifo-1(kc2)IV;ifb-2(kc14)II* was generated by crossing strain BJ142 with strain BJ309. Strains BJ411 *bbln-1(mib70)* and BJ412 *bbln-1(mib70);ifb-2(kc14)* were generated by crossing BOX415 with BJ309 resulting in strain BJ364 *erm-1(mib40[erm-1::AID::mCherry])I;Pelt-2::TIR::tagBFP-2::tbb-2-3’UTR (mib58[Pelt-2::TIR-1::tagBFP-2-Lox511::tbb-2-3’UTR])IV;C15C7.5(mib70[pC15C7.5::GFP1-3, C15C7.5 0-914del])X;ifb-2(kc14)II*, which was subsequently crossed with N2.

### Suppressor screen

10 cm diameter agar plates containing normal nematode growth medium (NGM) were prepared by autoclaving a solution containing 3 g NaCl (Carl Roth, Karlsruhe, Germany), 2.5 g Bacto peptone (BD BioSciences, Heidelberg, Germany) and 18.75 g Bacto agar (BD BioSciences) per 1 l and pouring the solution after addition of 1 mL cholesterol (5 mg/mL in ethanol), 0.5 mL 1 M CaCl_2_, 1 mL 1 M MgSO_4_ and 25 mL 1 M KH_2_PO_4_ (pH 6.0). After solidification, OP50 were placed on top of the agar and plates were incubated overnight at 37°C. They were then used for growing OLB18. Adult animals were bleached with a solution containing 170 µl 12% NaOCl, 100 µl 4M NaOH per 1 mL PBS (Biochrom, Berlin, Germany) and the resulting embryos were grown on new plates. The resulting synchronized L4 larvae were washed off with PBS and were then centrifuged in 15 ml Falcon tubes at 340xg for 2 min. 5 ml of the pelleted worms were mixed with 50 µl of 50 mM N-ethyl-N-nitrosourea (ENU; Sigma-Aldrich, Munich, Germany) and the suspension was incubated for 4 h at room temperature on a rotating shaker. Two washing steps with PBS followed before the mutagenized worms were resuspended in PBS and transferred onto 100 mutagenized L4 animals were placed per agar plate containing enriched NGM containing additionally 5 g Difco yeast extract (ThermoFisher Scientific, Massachusetts, USA) (total of 200 plates). The plates were then incubated at 20°C. It was ensured that the F2 generation was completely laid on these plates before food was used up. Then 1.5×1.5 cm pieces were cut out and placed on new plates, which were again incubated at 20°C until food was completely used up. After repeating these steps three times, the development of the mutant lines compared to OLB18 was monitored using a stereomicroscope. Criteria for suppressor activity were body size and developmental stage. If both were rescued, lines were subjected to further selection rounds. Only stable lines were subsequently examined by light microscopy for rescue of invaginations of the apical intestinal membrane. The mutant line, which met all three criteria best, was named strain BJ334 (*sma-5(n678)X;ifb-2(kc20)II*) and was backcrossed twice with OLB18 resulting in strain BJ355 (*sma-5(n678)X;ifb-2(kc20)II*). To determine the mutation of allele *kc20 ifb-2* DNA was amplified in three parts using three primer pairs outside the protein coding regions (oSMR66 (ggtgttggttttttaactgctg) and oSMR68 (acacacccatttcctccaga), oSMR67 (tctggaggaaatgggtgtgt), and oSMR70 (tccttgcggatacactctga), oSMR69 (tcggtagctataaccgcttca), and oSMR71 (caaggaaaggattcaatgggc)). The DNA was purified and analyzed by Sanger sequencing using the same primers as for amplification.

### CRISPR-Cas9

All alleles were made using homology-directed repair of CRISPR-Cas9-induced DNA double strand breaks, using microinjected Cas9 ribonucleoprotein complexes and linear ssDNA oligo repair templates, similar to the approach described by (Ghanta et al., 2021). We used 250-700 ng/µl Cas9, a Cas9/crRNA ratio of 3.0-4.5, and the pRF4 (*rol-6(su1006)*) co-injection marker (Mello et al., 1991). Repair templates included, when appropriate, silent mutations to prevent recutting of repaired loci by Cas9. To verify the edits, the insertion sites were PCR amplified and sequenced by Sanger sequencing.

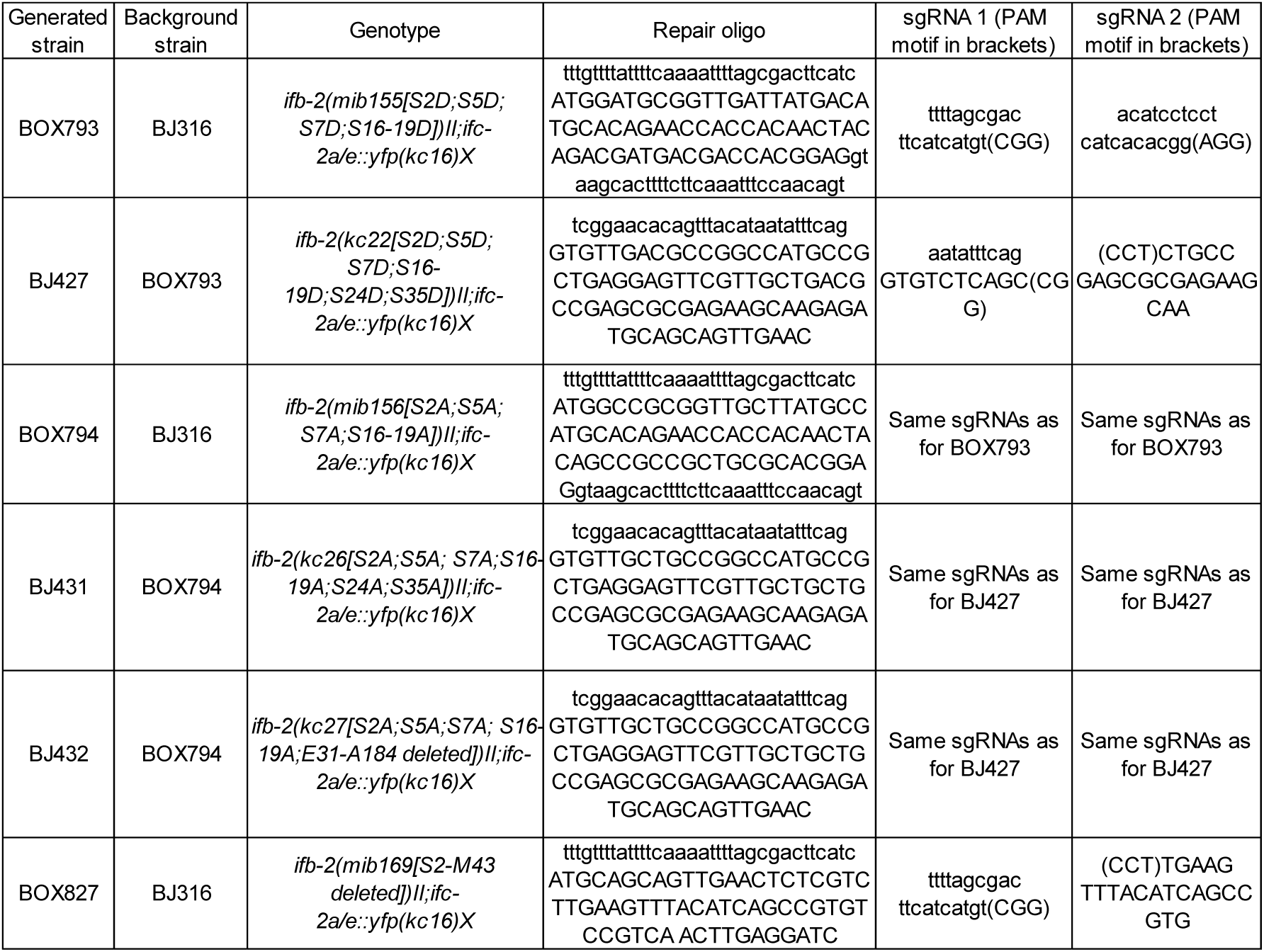

### Microscopy

Light microscopy was performed with a Zeiss (Jena, Germany) apotome in combination with a ZeissAxioCamMRm camera.

For electron microscopy worms were either chemically fixed (Fig. 5) or cryofixed at high pressure (Fig. 1):

#### Chemical fixation

Young adult animals (40 for each strain) were submerged in freshly prepared fixation solution containing 2.5% glutaraldehyde (using 25% (w/v) glutardialdehyde from Carl Roth), 1% (w/v) paraformaldehyde (using a stock solution with 0.4 g paraformaldehyde dissolved in 10 ml 0.1 M sucrose with 0.6 µl 10 N sodium hydroxide [all from Sigma-Aldrich]), 0.05 M cacodylate buffer (using dimethylarsinic acid sodium salt trihydrate from Merck and HCl for pH adjustment to 6.4-7.4) at room temperature in a glass staining block. Each worm was cut through at the anterior and posterior end with a scalpel. The fixation solution was exchanged two times with 1.5-2 h incubation at room temperature in between. After the third replacement of the fixation solution an overnight incubation followed at 4°C in a moist chamber. The samples were then transferred to 0.2 M cacodylate buffer and incubated for 3×10 min in this buffer with buffer changes in between. Samples were then incubated for 4 h in 0.1 M cacodylate buffer containing 1% (w/v) OsO_4_ (Paesel-Lorei, Frankfurt/Main, Germany) and 0.5% (w/v) K_3_[Fe(CN)_6_] (Carl Roth), followed by 3×10 min in 0.1 M cacodylate buffer. Sometimes samples were incubated overnight at 4°C in a moist chamber in 0.1 M cacodylate buffer. The following incubations ensued at room temperature: 3×10 min in 0.1 M maleic acid buffer (Sigma-Aldrich; pH 6), 2 h in the dark in 0.5% (w/v) uranyl acetate (EMS, Hatfield, PA, USA) dissolved in 0.05 N maleic acid buffer (pH 5.2), 3×5 min in 0.1 M maleic acid buffer, 3×5 min in double distilled water, 5 min in 20% ethanol, 5 min in 30% ethanol, 3×10 min in 50% ethanol, 2×15 min in 75% ethanol, 2×15 min in 95% ethanol, 3×10 min in 100% ethanol, 10 min in a 1:1 ethanol/acetone mixture and 2×10 min in acetone. Finally, samples were embedded in araldite (Agar scientific, Stansted, UK)). To this end samples were first placed overnight at 4°C in a 3:1 mixture of acetone/araldite with 1.5% (v/v) DMP30 [Agar scientific] followed by incubation steps at room temperature: 1 h in a 1:1 mixture of acetone/araldite with 1.5% DMP30, 2 h in a 1:3 mixture of acetone/araldite with 1.5% DMP30, and two days in araldite with 2% DMP30. The final polymerization was done at 60°C for two days in silicone molds with pre-polymerized araldite at the bottom. 75 nm sections were prepared using a Leica (Wetzlar, Germany) Reichert Ultracut S microtome. They were contrasted for 4 min in uranyl acetate and 3 min in lead citrate and finally imaged at 60 kV in a Zeiss EM 10.

#### High-pressure freezing

Young adult animals were transferred into a 100 μm deep membrane carrier containing 20% bovine serum albumin in M9 worm buffer (22 mM KH_2_PO_4_, 42 mM Na_2_HPO_4_, 86 mM NaCl, 1 mM MgSO_4_) and then high-pressure frozen in a Leica EM Pact high-pressure freeze. A minimum of five samples with 10-20 animals were frozen per experiment. Quick freeze substitution using 1% OsO_4_, 0.2% uranyl acetate in acetone followed by epoxy resin embedding was performed as previously described (McDonald and Webb, 2011). Subsequently, 50 nm thick sections of the embedded samples were prepared using a Leica UC6/FC6 ultramicrotome. These were contrasted for 10 min in 1% uranyl acetate in ethanol and Reynolds lead citrate and recorded at 100 kV on a Hitachi H-7600 transmission electron microscope (Tokyo, Japan).

### Immunoblotting

60 young adults of strains N2 and BJ432 were picked in 30 µl dH_2_O each and frozen at - 80°C. To disrupt the cuticle, the samples were then rapidly thawed on a heating block and three times sucked up and down through a 30G hypodermal syringe (BD Medical, Heidelberg, Germany). After addition of 7.5 µl 5x Laemmli loading buffer (15 ml stacking gel buffer [0.15 M Tris-Cl, 0.1% SDS, pH: 6,8], 12.5 ml glycerine, 2.5 ml β-mercaptoethanol, 2.5 g SDS, some bromophenol blue), the samples were incubated for 5 min at 90°C. Polypeptides were separated by electrophoresis in an 8% sodium dodecyl sulfate (SDS) polyacrylamide gel. Separated proteins were then transferred by wet tank-blotting (100 V for 60 min) onto a polyvinylidene difluoride (PVDF) membrane (Merck, Darmstadt, Germany). The membrane was blocked with Roti^®^-Block (2 h at room temperature; Carl Roth) and incubated overnight at 4°C with the primary antibody (mouse monoclonal anti-IFB-2 antibody MH33, 1:1000 dilution in Roti^®^-Block, Developmental Studies Hybridoma Bank, (Francis and Waterston, 1991)). The membrane was washed three times with TBST (20 mM tris(hydroxymethyl)-aminomethane, 0.15 M NaCl, 0.1% Tween 20 (v/v), pH 7.6) and then incubated with the secondary antibody (goat anti-mouse IgG antibodies coupled to horseradish peroxidase from DAKO at 1:5000 in Roti^®^-Block) for 1 h at room temperature. Chemiluminescence substrate AceGlow (VWR, Darmstadt, Germany, #730-1510) was detected by Fusion Solo (Vilber Lourmat, Eberhardzell, Germany).

### Analysis of larval development and progeny production

For the analysis of larval development, a defined number of isolated embryos were placed on a NGM plate with an OP50 bacterial lawn and incubated at 18°C. The number of adult stages was then determined daily, and the remaining larval stages were transferred to a new plate. For further in-depth analysis of the different stages of development, the individual age of each larvae was also determined. The larval arrest rate was calculated from the following quotient: number of animals that died as larvae divided by the total number of animals at the beginning of the experiment. Significance was calculated using the Chi square function of Excel (Microsoft, Redmont, WA). Time of development was calculated based on the time it took to complete development from the embryo (up to 24 cell stage) to the adult stage. The offspring of these animals was determined and is presented as average progeny per individual. The generated data are presented as mean value ± SD. Significance was calculated using the unpaired, two-tailed t-test function of GraphPad Prism 5.01 (GraphPad Software Inc., LaJolla, CA).

### Life span analysis

Embryos were isolated, transferred to NGM plates with OP50 bacterial lawns and then incubated at 18°C. The hatched animals were checked daily for vitality. Animals without reaction to mechanical stimulus, triggered by a platinum wire, were considered dead. Animals that could not be found were censored. In order to avoid mix up with the following generation, the animals were transferred to new plates at least every three days. Statistical analysis was performed using the survival function and the Gehan-Breslow-Wilcoxon test of GraphPad Prism 5.01.

### Stress assays

For the oxidative stress assay, NGM agar plates containing 200 mM methyl viologen dichloride hydrate (paraquat; Sigma-Aldrich) were prepared, stored at room temperature overnight and then inoculated with 50 μl of a 10 x concentrated OP50 overnight culture. After another day at room temperature plates were stored at 4°C and were used for a maximum of three days. L4 larvae were placed on the bacterial lawn and incubated at 18°C. Every hour the viability of the animals was checked by mechanically provoking them with a platinum wire. Vital animals showed an active response to this stimulus. Animals that did not respond were considered dead. In parallel, plates without paraquat were used as a control. For statistical analysis, the Gehan-Breslow-Wilcoxon test of GraphPad Prism 5.01 was used.

To perform the osmotic stress assay, plates containing 300 mM NaCl were prepared, stored overnight at room temperature, then inoculated with 300 μl of an OP50 overnight culture and incubated for another day at room temperature. Afterwards, L4 stages were placed on the bacterial lawn and plates were incubated overnight at 18°C. The animals were then washed in recovery buffer (M9 buffer with 150 mM NaCl) and transferred to normal NGM plates. After a further overnight incubation at 18°C, single worms were tested for viability as described above. Data shown correspond to mean value ± SD. Statistical calculations were performed using the Chi square function of Excel.

### RNAi and body length determination

RNAi by feeding was performed as described previously (Geisler et al., 2016) without supplement of tetracycline. In brief, RNAi plates were inoculated with 300 µl of overnight grown HT115 bacteria producing dsRNA targeted against *ifb-2*, *ifc-1/-2*, *ifd-1/-2* and *ifp-1* mRNA followed by overnight incubation at room temperature. L4 larvae were placed onto plates and incubated for 48 hours at 18°C. Subsequently, adult grown animals were transferred to new plates and incubated for 4 days at 18°C. Laid progeny was used for imaging and body length measurements. Animals were either imaged on the plate using a Nikon AZ100M stereoscope (Düsseldorf, Germany) equipped with a SONY Alpha 7R (Berlin, Germany) camera or mounted on agar slides using 10% levamisole as anaesthetic. Body length was determined using the segmented line tool in combination with the measurement function of Fiji (https://imagej.net/Fiji). Results are shown as mean value ± SD. Significance was calculated using the unpaired, two-tailed t-test function of GraphPad Prism 5.01.

RNAi-inducing bacteria were commercially available through the Vidal library (clone *ifb-2*, *ifc-2*, *ifd-2*, Source BioScience, Nottingham, UK), the Ahringer feeding library (clone *ifc-1*, Source BioScience, Nottingham, UK). RNAi clones for *ifd-1* and *ifp-1* were generated by subcloning 1 kb of the corresponding cDNA into a modified L4440 RNAi feeding vector, containing a linker with *AscI* and *NotI* restriction sites, analogous to the site of insertion. The following primers were used: oSMR57 (aggcgcgccTGACCACCATAGCCGAACTT), oSMR58 (agcggccgcTTTGAAGCCACCAACGTCTG), oSMR59 (aggcgcgccTCAAAACCGGGTTCTCGAGA), oSMR60 (agcggccgcTTCACTGCGGAGGTTGATCT). Vector identity was verified by DNA sequencing in each instance.

## Acknowledgments

We thank Barbara Bonn, Stephanie Brosig, Janis Moeller and Sabine Eisner for their commitment and excellent technical support and Adam Breitscheidel for thoughtful figure arrangement. We are particularly thankful to Christine Richardson and Martin Goldberg (Durham University, UK) for high-pressure freeze electron microscopy. Strains N2 and FK312 were obtained from the Caenorhabditis Genetics Centre (University of Minnesota, Minneapolis, MN). The MH33 monoclonal antibody was obtained from the Developmental Studies Hybridoma Bank developed under the auspices of the NICHD and maintained by the University of Iowa, Department of Biology, Iowa City, IA.

## Declarations

### Ethics approval

Not applicable.

### Consent to participate

All authors consented to participate.

### Consent for publication

All authors consented to publication.

### Availability of data and material

The authors confirm that all relevant data are included in the article.

### Authors’ contributions

Conceptualization: F.G., R.E.L.; Methodology: F.G.; Validation: F.G.; Formal analysis: F.G.; Investigation: F.G., S.R.; Resources: R.E.L.; Data curation: F.G.; Writing - original draft: F.G., R.E.L.; Visualization: F.G.; Supervision: F.G., M.B., R.E.L.; Project administration: R.E.L.; Funding acquisition: F. G., M.B., R.E.L.

## Supplementary Figures

**Figure S1.**
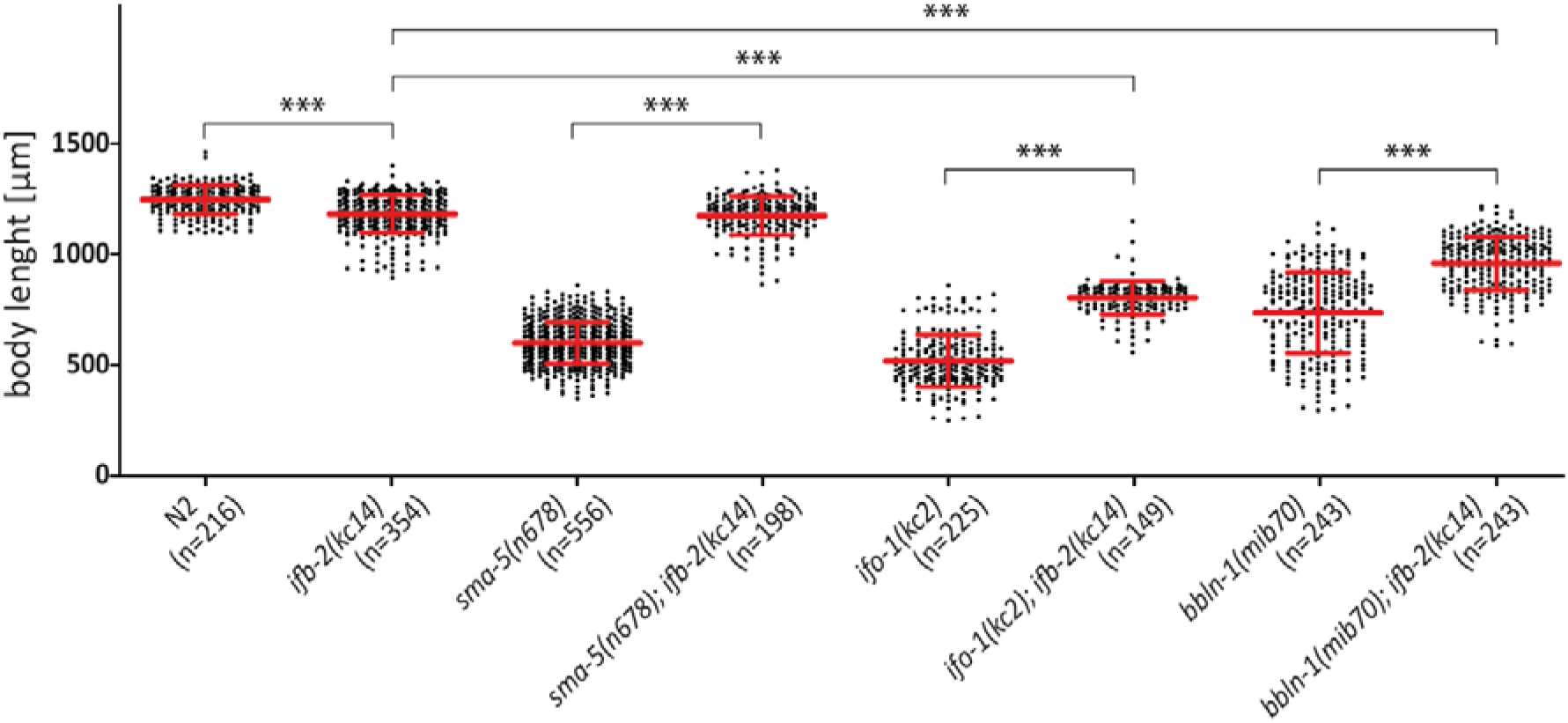
Depletion of IFB-2 rescues growth defects in *sma-5(n678)*, *ifo-1(kc2)* and *bbln-1(mib70)* loss of function mutants. The scatter dot blot shows a comparison of body lengths 4 days after egg laying in N2, *ifb-2(kc14)*, *sma-5(n678)*, *sma-5(n678);ifb-2(kc14)*, *ifo-1(kc2)*, *ifo-1(kc2);ifb-2(kc14)*, *bbln-1(mib70)* and *bbln-1(mib70);ifb-2(kc14)*. Note that loss of IFB-2 rescues the body length phenotype of *sma-5(n678)* to the level of *ifb-2(kc14)* single mutants (N2: 1252±65.59 µm; *ifb-2(kc14)*: 1187±84.85 µm; *sma-5(n678)*: 608.2±91.97 µm; *sma-5(n678);ifb-2(kc14)*: 1179±87.57 µm; *** p<0.0001). Rescue of body length is also observed in *ifo-1(kc2)* and *bbln-1(mib70)* albeit less efficiently and to different degrees (*ifo-1(kc2)*: 526.7.7±117.2 µm; *ifo-1(kc2);ifb-2(kc14)*: 809.6.3±75.2 µm; *** p<0.0001; *bbln-1(mib70)*: 742.7±181.3 µm; *bbln-1(mib70);ifb-2(kc14)*: 964.3±120.1 µm; *** p<0.0001).

**Figure S2.**
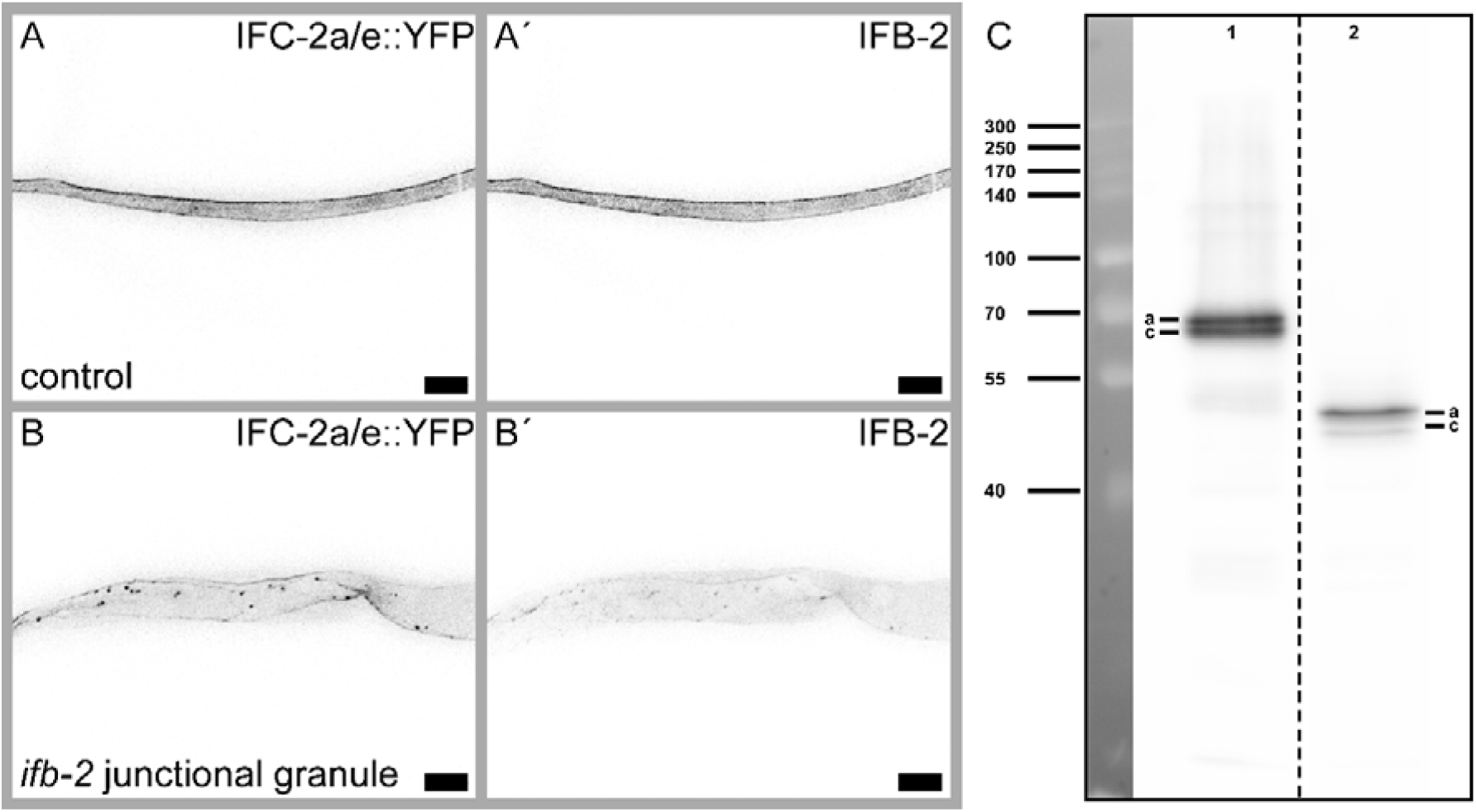
*ifb-2(kc27)* animals show IFC-2 positive junctional granules in concert with an overall reduced IFB-2/IFC-2-containing network. **(A-B**’) The fluorescence micrographs show IFC-2a/e::YFP **(A, B)** and corresponding IFB-2 immunostainings **(A’, B’)** of isolated control **(A-A’)** and *ifb-2(kc27[S2A;S5A;S7A;S16-19A;E31-A184 deleted])II* intestines. Note the reduced IFC-2a/e::YFP in the apical cytoplasm with additional IFC-2-positive junctional granules. In contrast, IFB-2 is also diffusely localized in the cytoplasm. **(C)** Immunoblot detecting IFB-2 isoforms a and c of control (lane 1) and *ifb-2(kc27)* animals (lane 2). The IFB-2 deletion in *ifb-2(kc27)* results in a truncated protein of about 50 kDa.

## References

Al-Hashimi, H., Hall, D. H., Ackley, B. D., Lundquist, E. A. & Buechner, M. 2018. Tubular Excretory Canal Structure Depends on Intermediate Filaments EXC-2 and IFA-4 in Caenorhabditis elegans. Genetics, 210, 637–652.

Bossinger, O., Fukushige, T., Claeys, M., Borgonie, G. & Mcghee, J. D. 2004. The apical disposition of the Caenorhabditis elegans intestinal terminal web is maintained by LET-413. Dev Biol, 268, 448–56.

Bott, C. J. & Winckler, B. 2020. Intermediate filaments in developing neurons: Beyond structure. Cytoskeleton (Hoboken), 77, 110–128.

Carberry, K., Wiesenfahrt, T., Geisler, F., Stocker, S., Gerhardus, H., Uberbach, D., Davis, W., Jorgensen, E., Leube, R. E. & Bossinger, O. 2012. The novel intestinal filament organizer IFO-1 contributes to epithelial integrity in concert with ERM-1 and DLG-1. Development, 139, 1851–62.

Carberry, K., Wiesenfahrt, T., Windoffer, R., Bossinger, O. & Leube, R. E. 2009. Intermediate filaments in Caenorhabditis elegans. Cell Motil Cytoskeleton, 66, 852–64.

Chamcheu, J. C., Siddiqui, I. A., Syed, D. N., Adhami, V. M., Liovic, M. & Mukhtar, H. 2011. Keratin gene mutations in disorders of human skin and its appendages. Arch Biochem Biophys, 508, 123–37.

Clemen, C. S., Herrmann, H., Strelkov, S. V. & Schroder, R. 2013. Desminopathies: pathology and mechanisms. Acta Neuropathol, 125, 47–75.

Coch, R., Geisler, F., Annibal, A., Antebi, A. & E. Leube, R. 2020. Identification of A Novel Link between the Intermediate Filament Organizer IFO-1 and Cholesterol Metabolism in the Caenorhabditis elegans Intestine. International Journal of Molecular Sciences, 21, 8219.

Coch, R. A. & Leube, R. E. 2016. Intermediate Filaments and Polarization in the Intestinal Epithelium. Cells, 5.

Coulombe, P. A., Kerns, M. L. & Fuchs, E. 2009. Epidermolysis bullosa simplex: a paradigm for disorders of tissue fragility. J Clin Invest, 119, 1784–93.

Coulombe, P. A. & Omary, M. B. 2002. ’Hard’ and ’soft’ principles defining the structure, function and regulation of keratin intermediate filaments. Curr Opin Cell Biol, 14, 110–22.

D’alessandro, M., Russell, D., Morley, S. M., Davies, A. M. & Lane, E. B. 2002. Keratin mutations of epidermolysis bullosa simplex alter the kinetics of stress response to osmotic shock. J Cell Sci, 115, 4341–51.

Didonna, A. & Opal, P. 2019. The role of neurofilament aggregation in neurodegeneration: lessons from rare inherited neurological disorders. Mol Neurodegener, 14, 19.

Estes, K. A., Szumowski, S. C. & Troemel, E. R. 2011. Non-lytic, actin-based exit of intracellular parasites from C. elegans intestinal cells. PLoS Pathog, 7, e1002227.

Etienne-Manneville, S. 2018. Cytoplasmic Intermediate Filaments in Cell Biology. Annu Rev Cell Dev Biol, 34, 1–28.

Francis, R. & Waterston, R. H. 1991. Muscle cell attachment in Caenorhabditis elegans. J Cell Biol, 114, 465–79.

Geisler, F., Coch, R. A., Richardson, C., Goldberg, M., Bevilacqua, C., Prevedel, R. & Leube, R. E. 2020. Intestinal intermediate filament polypeptides in C. elegans: Common and isotype-specific contributions to intestinal ultrastructure and function. Sci Rep, 10, 3142.

Geisler, F., Coch, R. A., Richardson, C., Goldberg, M., Denecke, B., Bossinger, O. & Leube, R. E. 2019. The intestinal intermediate filament network responds to and protects against microbial insults and toxins. Development, 146.

Geisler, F., Gerhardus, H., Carberry, K., Davis, W., Jorgensen, E., Richardson, C., Bossinger, O. & Leube, R. E. 2016. A novel function for the MAP kinase SMA-5 in intestinal tube stability. Mol Biol Cell, 27, 3855–3868.

Geisler, F. & Leube, R. E. 2016. Epithelial Intermediate Filaments: Guardians against Microbial Infection? Cells, 5.

Gentil, B. J., Tibshirani, M. & Durham, H. D. 2015. Neurofilament dynamics and involvement in neurological disorders. Cell Tissue Res, 360, 609–20.

Ghanta, K. S., Chen, Z., Mir, A., Dokshin, G. A., Krishnamurthy, P. M., Yoon, Y., Gallant, J., Xu, P., Zhang, X. O., Ozturk, A. R., Shin, M., Idrizi, F., Liu, P., Gneid, H., Edraki, A., Lawson, N. D., Rivera-Perez, J. A., Sontheimer, E. J., Watts, J. K. & Mello, C. C. 2021. 5’-Modifications improve potency and efficacy of DNA donors for precision genome editing. Elife, 10.

Hatzfeld, M. & Burba, M. 1994. Function of type I and type II keratin head domains: their role in dimer, tetramer and filament formation. J Cell Sci, 107 (Pt 7), 1959–72.

Hüsken, K., Wiesenfahrt, T., Abraham, C., Windoffer, R., Bossinger, O. & Leube, R. E. 2008. Maintenance of the intestinal tube in Caenorhabditis elegans: the role of the intermediate filament protein IFC-2. Differentiation, 76, 881–96.

Jacob, J. T., Coulombe, P. A., Kwan, R. & Omary, M. B. 2018. Types I and II Keratin Intermediate Filaments. Cold Spring Harb Perspect Biol, 10.

Jahnel, O., Hoffmann, B., Merkel, R., Bossinger, O. & Leube, R. E. 2016. Mechanical Probing of the Intermediate Filament-Rich Caenorhabditis Elegans Intestine. Methods Enzymol, 568, 681–706.

Kao, C. Y., Los, F. C., Huffman, D. L., Wachi, S., Kloft, N., Husmann, M., Karabrahimi, V., Schwartz, J. L., Bellier, A., Ha, C., Sagong, Y., Fan, H., Ghosh, P., Hsieh, M., Hsu, C. S., Chen, L. & Aroian, R. V. 2011. Global functional analyses of cellular responses to pore-forming toxins. PLoS Pathog, 7, e1001314.

Karabinos, A., Schmidt, H., Harborth, J., Schnabel, R. & Weber, K. 2001. Essential roles for four cytoplasmic intermediate filament proteins in Caenorhabditis elegans development. Proc Natl Acad Sci U S A, 98, 7863–8.

Karabinos, A., Schunemann, J. & Parry, D. A. 2017. Assembly studies of six intestinal intermediate filament (IF) proteins B2, C1, C2, D1, D2, and E1 in the nematode C. elegans. Cytoskeleton (Hoboken), 74, 107–113.

Karabinos, A., Schunemann, J. & Weber, K. 2004. Most genes encoding cytoplasmic intermediate filament (IF) proteins of the nematode Caenorhabditis elegans are required in late embryogenesis. Eur J Cell Biol, 83, 457–68.

Koyuncu, S., Loureiro, R., Lee, H. J., Wagle, P., Krueger, M. & Vilchez, D. 2021. Rewiring of the ubiquitinated proteome determines ageing in C. elegans. Nature, 596, 285–290.

Leube, R. E. & Schwarz, N. 2016. Intermediate Filaments. In: Bradshaw, R. A. & Stahl, P. D. (eds.) Encyclopedia of Cell Biology. Waltham, MA: Academic Press.

Magin, T. M., Vijayaraj, P. & Leube, R. E. 2007. Structural and regulatory functions of keratins. Exp Cell Res, 313, 2021–32.

Margiotta, A. & Bucci, C. 2016. Role of Intermediate Filaments in Vesicular Traffic. Cells, 5.

Mcdonald, K. L. & Webb, R. I. 2011. Freeze substitution in 3 hours or less. J Microsc, 243, 227–33.

Mcghee, J. D. 2007. The C. elegans intestine (March 27, 2007), WormBook, ed. The C. elegans Research Community, WormBook, doi/10.1895/wormbook.1.133.1, http://www.wormbook.org.

Mello, C. C., Kramer, J. M., Stinchcomb, D. & Ambros, V. 1991. Efficient gene transfer in C.elegans: extrachromosomal maintenance and integration of transforming sequences. EMBO J, 10, 3959–70.

Omary, M. B., Ku, N. O., Tao, G. Z., Toivola, D. M. & Liao, J. 2006. “Heads and tails“ of intermediate filament phosphorylation: multiple sites and functional insights. Trends Biochem Sci, 31, 383–94.

Pekny, M. & Lane, E. B. 2007. Intermediate filaments and stress. Exp Cell Res, 313, 2244–54.

Remmelzwaal, S., Geisler, F., Stucchi, R., van der Horst, S., Pasolli, M., Kroll, J. R., Jarosinska, O. D., Akhmanova, A., Richardson, C. A., Altelaar, M., Leube, R. E., Ramalho, J. J. & Boxem, M. 2021. BBLN-1 is essential for intermediate filament organization and apical membrane morphology. Curr Biol, 31, 2334–2346 e9.

Sawant, M. S. & Leube, R. E. 2017. Consequences of Keratin Phosphorylation for Cytoskeletal Organization and Epithelial Functions. Int Rev Cell Mol Biol, 330, 171–225.

Snider, N. T. & Omary, M. B. 2014. Post-translational modifications of intermediate filament proteins: mechanisms and functions. Nat Rev Mol Cell Biol, 15, 163–77.

Stutz, K., Kaech, A., Aebi, M., Kunzler, M. & Hengartner, M. O. 2015. Disruption of the C. elegans Intestinal Brush Border by the Fungal Lectin CCL2 Phenocopies Dietary Lectin Toxicity in Mammals. PLoS One, 10, e0129381.

Toivola, D. M., Strnad, P., Habtezion, A. & Omary, M. B. 2010. Intermediate filaments take the heat as stress proteins. Trends Cell Biol, 20, 79–91.

Watanabe, N., Nagamatsu, Y., Gengyo-Ando, K., Mitani, S. & Ohshima, Y. 2005. Control of body size by SMA-5, a homolog of MAP kinase BMK1/ERK5, in C. elegans. Development, 132, 3175–84.

Yoon, S. & Leube, R. E. 2019. Keratin intermediate filaments: intermediaries of epithelial cell migration. Essays Biochem, 63, 521–533.

Yoshida, T. & Nakagawa, M. 2012. Clinical aspects and pathology of Alexander disease, and morphological and functional alteration of astrocytes induced by GFAP mutation. Neuropathology, 32, 440–6.

Zhou, X., Lin, Y., Kato, M., Mori, E., Liszczak, G., Sutherland, L., Sysoev, V. O., Murray, D. T., Tycko, R. & Mcknight, S. L. 2021. Transiently structured head domains control intermediate filament assembly. Proc Natl Acad Sci U S A, 118.

